# An updated nomenclature for plant ribosomal protein genes

**DOI:** 10.1101/2022.07.06.498169

**Authors:** M. Regina Scarpin, Michael Busche, Ryan E. Martinez, Lisa C. Harper, Leonore Reiser, Ting Lan, Wei Xiong, Beixin Mo, Guiliang Tang, Xuemei Chen, Julia Bailey-Serres, Karen Browning, Jacob O. Brunkard

**Affiliations:** Laboratory of Genetics, University of Wisconsin – Madison, Madison, WI 53706, USA; Department of Plant and Microbial Biology, University of California – Berkeley, Berkeley, CA 94720, USA; Plant Gene Expression Center, USDA Agricultural Research Service, Albany, CA 94710, USA; Corn Insects and Crop Genetics Research Unit, USDA Agricultural Research Service, Ames, IA 50011, USA; The Arabidopsis Information Resource, Phoenix Bioinformatics, Fremont, CA 94538, USA; Guangdong Provincial Key Laboratory for Plant Epigenetics, Longhua Bioindustry and Innovation Research Institute, College of Life Sciences and Oceanography, Shenzhen University, Shenzhen, 518060 China; Key Laboratory of Optoelectronic Devices and Systems of Ministry of Education and Guangdong Province, College of Optoelectronic Engineering, Shenzhen University, Shenzhen, 518060 China; Department of Biological Sciences, Life Science and Technology Institute, Michigan Technological University, Houghton, MI 49931, USA; Department of Botany and Plant Sciences and Center for Plant Cell Biology, Institute of Integrative Genome Biology, University of California – Riverside, CA 92521, USA; Department of Molecular Biosciences, Center for Systems and Synthetic Biology, University of Texas, Austin, TX 78712, USA

## Abstract

Ban *et al*. (2014) proposed a nomenclature for ribosomal proteins (r-proteins) that reflects the current understanding of ribosomal protein evolution. In the past few years, this nomenclature has been widely adopted among biomedical researchers and microbiologists. This homology-based r-protein nomenclature has not been as widely adopted among plant biologists, however, presumably because r-protein nomenclature is much more complicated in plants due to gene duplication. Here, we propose compatible upgrades to the homology-guided nomenclature proposed by Ban *et al*. (2014) so that this naming system can be adopted for widespread use in the plant biology community. We note that Lan *et al*. (2022) recently proposed updated nomenclature for plant cytosolic ribosomal proteins, focused on Arabidopsis and rice. The nomenclature outlined here is an extension of that proposed by Lan *et al*. (2022), expanding to include organellar ribosomes and additional species, with the intent that this nomenclature can serve as a template to guide future plant genome annotations. A more detailed comparison highlighting how this naming system builds on the Ban *et al*. (2014) and Lan *et al*. (2022) nomenclatures is offered below.

At this time, we request **community feedback** on this proposed nomenclature so that the naming system ultimately chosen represents a broad consensus. Feedback can be communicated to the this working group at before July 25^th^, 2022. Coauthors of this letter and anyone in the scientific community expressing significant interest will then discuss this feedback as a group, reach a consensus agreement, and communicate the updated nomenclature rules through a letter to the editor (expected to be published at *The Plant Cell*) and the databases at TAIR and MaizeGDB.

---

Across all living organisms, ribosomes are large macromolecular complexes that synthesize proteins by translating messenger RNA codes into amino acid sequences. Structurally, ribosomes are composed of ~50-80 ribosomal proteins (r-proteins) and 3 or 4 ribosomal RNAs (rRNAs). Over the past 4 billion years, ribosomes have evolved some differences in rRNA and r-protein composition, with certain subunits specific to bacteria, archaea and eukaryotes, plastids, or mitochondria, although many subunits are universally conserved with clear homology across all of life. Historically, the nomenclature of r-proteins was different in each species investigated, based on biochemical properties of these r-proteins, which were numbered in the order that they were separated by electrophoresis and/or chromatography (for example, see (Wittmann *et al*. 1971)), rather than named for homology of structure or function. The different naming systems fostered confusion for researchers, especially scientists not directly investigating ribosome biology, and hindered computational efforts to collate information on homologous r-proteins. Ban *et al*. (2014) proposed their r-protein nomenclature rectify these issues.

In most lineages other than plants, r-proteins are encoded by single-copy genes (Steel and Jacobson 1986; Uechi et al. 2001). There are some small exceptions, of course; for example, bacterial genomes often include a couple of duplicated r-protein genes (Yutin et al. 2012), including *E. coli*, which has two copies of *bL31* and two copies of *bL36* (Makarova et al. 2001). *S. cerevisiae*, a descendent of a recent whole-genome duplication event, has two homeologous copies of many r-protein genes (Mager et al. 1997). Plant genomes, in contrast, almost always encode multiple paralogous copies of r-protein genes. For example, in *Arabidopsis thaliana*, every cytosolic r-protein is encoded by at least two paralogues, and several are encoded by five or six paralogues (Barakat et al. 2001; Salih et al. 2020; Lan et al. 2022). Moreover, plants also encode an additional two sets of r-proteins that localize in mitochondria or plastids to translate the organellar genomes. In sum, the Arabidopsis genome includes nearly 400 genes that encode r-proteins, about four-times more than the ~100 genes that encode r-proteins in mammals.

In consultation with The Arabidopsis Information Resource (TAIR), Maize Genetics and Genomics Database (MaizeGDB), and colleagues in the plant ribosome biology field, we have proposed new names for all of the r-proteins encoded by the Arabidopsis, tomato, maize, and rice genomes, which we intend will serve as a template to guide future plant genome annotations (Supplemental Dataset 1). We expect that this new nomenclature will enable greater communication with the wider audience of molecular biologists studying ribosomes and translation beyond plant biology.

The r-protein nomenclature established by Ban *et al*. (2014) begins with a lowercase letter indicating whether the r-protein is specific to bacteria (with the letter “b”), archaea and eukaryotes (with the letter “e”), or all domains of life (with the letter “u” for “universal”). This is followed by either L or S to indicate whether the protein is a subunit of the large or small ribosomal subunit, respectively, and then by a number to specify the r-protein identity (Fig. 1A). Cytosolic r-proteins have no suffix, whereas organelle-targeted r-protein names conclude with a suffix to indicate that they are targeted to mitochondria (with the letter “m”) or plastids (with the letter “c”, for “chloroplast”) (Bieri et al. 2017).

**Figure 1.**
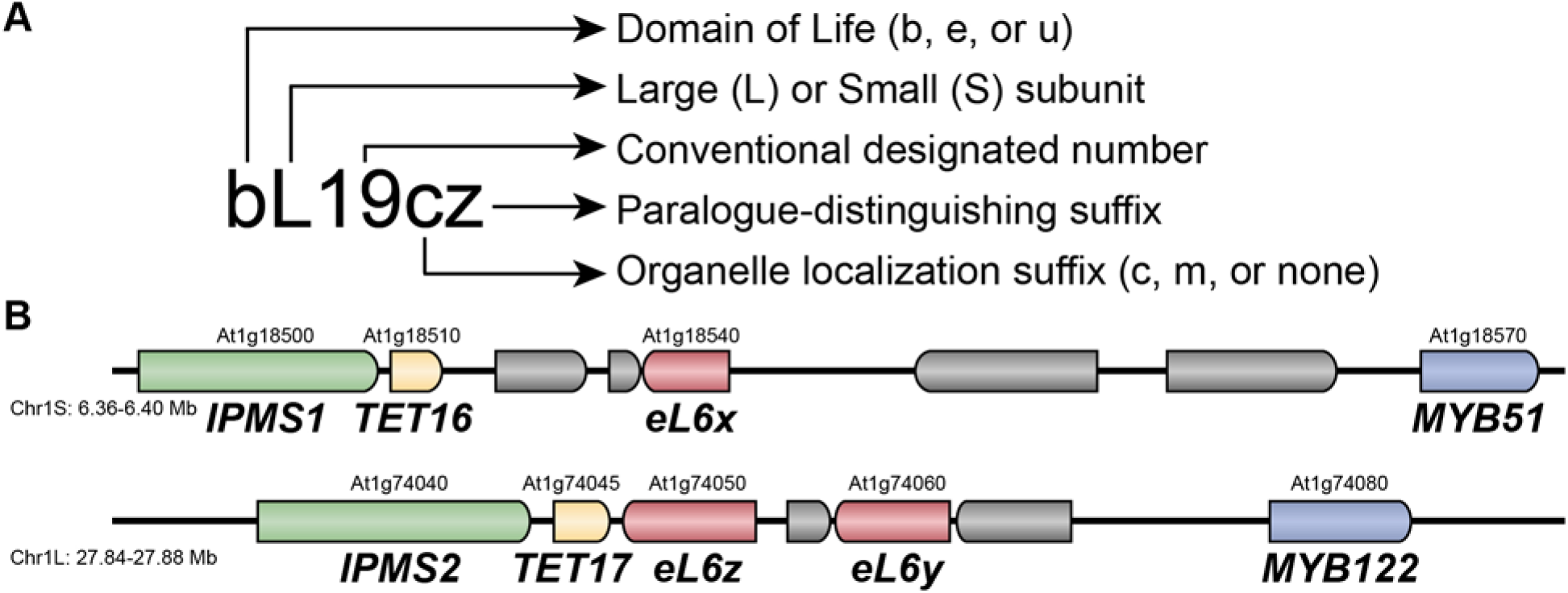
(A) The proposed r-protein nomenclature follows standard rules across all domains of life to indicate homology of ribosomal subunits. The first letter indicates where the r-protein is specific to bacterial genomes (b), archaean/eukaryotic genomes (e), or universal across genomes (u). The second letter indicates whether the r-protein is associated with the large 60S (L) or small 40S (S) subunit. The subunit number is based on consensus convention across model species as previously established (Ban *et al*. 2014). R-proteins that localize to plastids (c) or mitochondria (m) are indicated with a suffix. The final suffix is used to distinguish paralogues that encode homologous r-proteins within a genome. (B) Representative example of r-protein paralogy in the *Arabidopsis thaliana* genome. *eL6x* is a homeologue of two tandemly duplicated paralogues, *eL6z* and *eL6y*. Neighboring homeologous genes and chromosomal locations are indicated to demonstrate synteny among these r-protein genes.

Where feasible, the new r-protein names retain their traditional numbers—for example, archaeal/eukaryotic RPS6 is now eS6. Bacterial RPS6 is not homologous to eukaryotic RPS6, however, which previously caused some confusion; now, bacterial RPS6 is bS6, to indicate that it is not related to any archaeal/eukaryotic r-protein. Conversely, uS8 is now the universal name for bacterial r-protein S8, yeast r-protein S22, and human r-protein S15A, which all had different names despite their homology. Plant r-proteins occasionally have their own names, as well; for example, uL3, which was previously called L3 in bacteria, humans, and yeast, is called RIBOSOMAL PROTEIN 1 (RP1) in Arabidopsis. Many Arabidopsis cytosolic r-proteins were first characterized from genetic screens for developmental defects, and the genes encoding these proteins were first named according to their mutant phenotypes, such as *apiculata*, *embryo defective*, *evershed*, *hapless*, *oligocellula*, *piggyback*, *pointed first leaves*, *short valve*, and *suppressor of acaulis*. Bifunctional r-proteins, such as eL40, which is proteolytically cleaved during ribosome assembly to separate the mature eL40 protein and its fused ubiquitin domain, are occasionally named not for the r-protein subunit, but for ubiquitin (in Arabidopsis, eL40 is called UBIQUITIN EXTENSION PROTEIN or UBQ, for example). These examples clearly illustrate the need for the new, unifying nomenclature for r-proteins in plant genomes.

The greatest challenge in adopting this new nomenclature for plant biology is how to best indicate paralogy of r-proteins (Fig. 1B). In the simplest cases, there are only two paralogues, which could be designated with a single letter in alphabetical order, e.g., eS6a and eS6b. But in many cases, there are at least three paralogues, which is problematic because the plastid-targeted proteins are designated with a “c” (Bieri et al. 2017). In Arabidopsis, about 20 cytosolic r-proteins would end with a “c” and be confused with the homologous plastid-targeted r-proteins that would also end with a “c”. There are many possible solutions to this problem; the most straightforward options are (1) to switch from a “c” designating chloroplast-targeted to a “p” designating plastid-targeted, (2) to add a hyphen separating the paralogue designation from the protein name, (3) to distinguish between majuscule (uppercase) and miniscule (lowercase) lettering, such that “C” indicates a third paralogue but “c” indicates plastid localization, or (4) to start from the end of the alphabet, naming paralogues, e.g., uL15z, uL15y, uL15x. We tentatively prefer the last option for several reasons. First, there is already literature on chloroplast ribosomes using the “c” to indicate plastid-targeted r-proteins. Second, “p” is used as a suffix in many nomenclatures to distinguish proteins from nucleic acids (e.g., Tor1p is the protein encoded by the gene *tor1* in fission yeast) or to designate protein phosphorylation (e.g., rpS6P is phosphorylated eS6). Third, hyphens are typically used in plant nomenclatures to indicate alleles, so naming genes *eS6-a* and *eS6-b* could give the false impression that these are two alleles of a single gene, rather than paralogues. Finally, relying on uppercase versus lowercase letters would require that database curators, computational biologists annotating new genomes, journal editors, and ribosome biologists working outside plant biology all pay strict attention to a slight typographical difference, whereas starting from the end of the alphabet avoids any potential confusion. Although we present that option here (Supplemental Dataset 1), each alternative has merits that can be considered by the community.

We have provided a provisional table of r-protein names for Arabidopsis, tomato, maize, and rice for the plant biology community to consider, alongside their historical names in Arabidopsis and their names as recently proposed by Lan *et al*. (2022) (Supplemental Dataset 1). Note that the Lan *et al*. (2022) nomenclature differs primarily in how paralogues are named, which is a result of the exclusive focus of that nomenclature on cytosolic ribosomes. The new nomenclature will be added to public databases, including TAIR, MaizeGDB, the Plant Cytoplasmic Ribosomal Proteins database (PlantCRP.cn), and any other databases recommended by the community. Previous names will be retained at these databases as a reference, and in publications, systematic identifiers (e.g., the Arabidopsis Genome Initiative (AGI) locus name) should always be used alongside the updated r-protein names. We strongly encourage researchers to adopt the new nomenclature once it is adopted to facilitate communication with researchers outside the plant community and increase the impact of our community’s work on ribosome biology.

## Methods

### Identifying Arabidopsis r-proteins and their subcellular localization

Arabidopsis r-proteins were identified by previous curation of gene models in Araport11 and in TAIR, by additional curation of cytosolic r-proteins from Lan *et al*. (2022), and proteomic analyses of ribosomal proteins by Salih *et al*. (2020). SUBA4 (Hooper et al. 2017) was used to further validate the subcellular localization of all cytosolic and organellar r-proteins.

### Identifying orthologues of r-proteins in tomato, rice, and maize

Orthologues of Arabidopsis r-proteins were found in tomato (*Solanum lycopersicum*), rice (*Oryza sativa*), and maize (*Zea mays*) using eggNOG 5.0 (Huerta-Cepas et al. 2019), PANTHER 14 (Mi et al. 2019), and Ensembl (Cunningham et al. 2022). For the purpose of this proposal, annotations were made conservatively, excluding annotated genes that could not be placed into an r-protein superfamily unambiguously or for which no expression data (transcriptomic or proteomic) is publicly available.

## Author Contributions and Acknowledgments

This work was supported by NIH DP5-OD023072 to J.O.B.

## Notes

### Competing Interest Statement

The authors have declared no competing interest.

